# An integrated map of cell type-specific gene expression in pancreatic islets

**DOI:** 10.1101/2023.02.03.526994

**Authors:** Ruth M Elgamal, Parul Kudtarkar, Rebecca L Melton, Hannah M Mummey, Paola Benaglio, Mei-Lin Okino, Kyle J Gaulton

## Abstract

Pancreatic islets are comprised of multiple endocrine cell types that produce hormones required for glucose homeostasis, and islet dysfunction is a major factor in the development of type 1 and type 2 diabetes (T1D, T2D). Numerous studies have generated gene expression profiles in individual islet cell types using single cell assays. However, there is no canonical reference of gene expression in islet cell types in both health and disease that is also easily accessible for researchers to access, query, and use in bioinformatics pipelines. Here we present an integrated reference map of islet cell type-specific gene expression from 192,203 cells derived from single cell RNA-seq assays of 65 non-diabetic, T1D autoantibody positive (Aab+), T1D, and T2D donors from the Human Pancreas Analysis Program. We identified 10 endocrine and non-endocrine cell types as well as sub-populations of several cell types, and defined sets of marker genes for each cell type and sub-population. We tested for differential expression within each cell type in T1D Aab+, T1D, and T2D states, and identified 1,701 genes with significant changes in expression in any cell type. Most changes were observed in beta cells in T1D, and, by comparison, there were almost no genes with changes in T1D Aab+. To facilitate user interaction with this reference, we provide the data using several single cell visualization and reference mapping tools as well as open-access analytical pipelines used to create this reference. The results will serve as a valuable resource to investigators studying islet biology and diabetes.

## Introduction

The islets of Langerhans in the pancreas are clusters of endocrine cells including alpha, beta, delta, and gamma cell types which each produce hormones that regulate blood glucose levels (1). Dysfunction of beta cells is one of the major pathologies of both type 1 and type 2 diabetes, which collectively affect over 500 million individuals worldwide (2,3). Other cell types in the microenvironment around islets also contribute to the modulation of islet function and diabetes risk such as endothelial and immune cells (4,5). The regulation of gene activity establishes the identity of specific cell types as well as changes in response to environmental stimuli and disease states, and gene activity can be measured by sequence-based expression profiling (6). Understanding the gene expression profiles of islet cell types can therefore provide insight into their function and can also reveal how cells are altered in diabetes.

Single cell technologies enable profiling the expression levels of genes in individual cells, which can then be used to define the gene regulatory profiles of specific cell types (7,8). Numerous studies have assayed gene expression in individual islet cells using single cell techniques (9–13). These studies have defined gene expression profiles of endocrine and non-endocrine cell types in the pancreas, heterogeneous sub-populations of cells representing cellular states within cell types, and changes in disease states including T1D and T2D. A caveat to these studies is that they have been performed using limited sample numbers, and in some cases limited cell numbers, and there has been inconsistency in the results across studies particularly when describing heterogeneity and changes in disease (14,15). In addition, the results of islet single cell studies are not often made easily accessible to researchers, particularly those that are not experts in single cell data analysis, to visualize and query the expression of a gene in each cell type or changes in cell type expression in disease.

The Human Pancreas Analysis Program (HPAP) was developed to comprehensively collect and profile pancreatic islet tissue from human donors to understand the pathogenesis of T1D and T2D (16,17). The data generated by HPAP for each donor includes bulk and single cell RNA-seq and ATAC-seq data as well as histology, genotyping, cellular phenotyping, and other data types. Data generated by HPAP are made freely available to researchers via a web portal PANC-DB (https://hpap.pmacs.upenn.edu/) where the raw sequence files from each individual donor can be directly downloaded (16). The rich set of raw islet donor data provided by this resource can then be used by researchers to create integrated resources which make HPAP accessible to the wider community studying islet biology and diabetes to develop testable hypotheses.

In this study we created a reference map of gene expression in pancreatic islet cell types using single cell RNA-seq data from 65 donors available in HPAP. Using this reference map, we created several additional resources including (i) marker gene lists for every islet cell type and subpopulation, (ii) normalized expression levels of genes in each islet cell type, and (iii) changes in gene expression in T1D, T1D autoantibody positive (Aab+), and T2D states in each islet cell type. We host these data in several interactive applications to enable researchers to visualize, query, and analyze this reference. Finally, we provide open-source analytical pipelines used to create the reference map and annotations. These resources are available at www.isletgenomics.org.

## Results

### Reference map of single cell expression in islets

We downloaded single cell RNA-seq (scRNA-seq) data from 67 donors available in PANC-DB (**Supplementary Table 1**). After pre-filtering of barcodes for each sample based on >500 expressed genes, we excluded two samples which had lower average expressed genes per cell. With the remaining 65 samples, we performed processing and clustering using a custom pipeline. In brief, this pipeline consists of ambient RNA background correction, dimension reduction of log normalized counts, batch correction, Leiden clustering, and post-clustering doublet removal. (**see Methods** for more detail). The resulting map had 192,203 cells which mapped to 14 distinct clusters (**Figure 1A**). In the final map, on average, samples had 2,957 cells with 16,908 unique molecular identifiers (UMI) per cell. Clusters were broadly consistent across samples, and no clusters were preferentially represented by a small number of samples (**Supplementary Figure 1**). We also observed little evidence for residual batch effects in the clusters driven by donor or other variables (**Figure 1B**).

**Figure 1.**
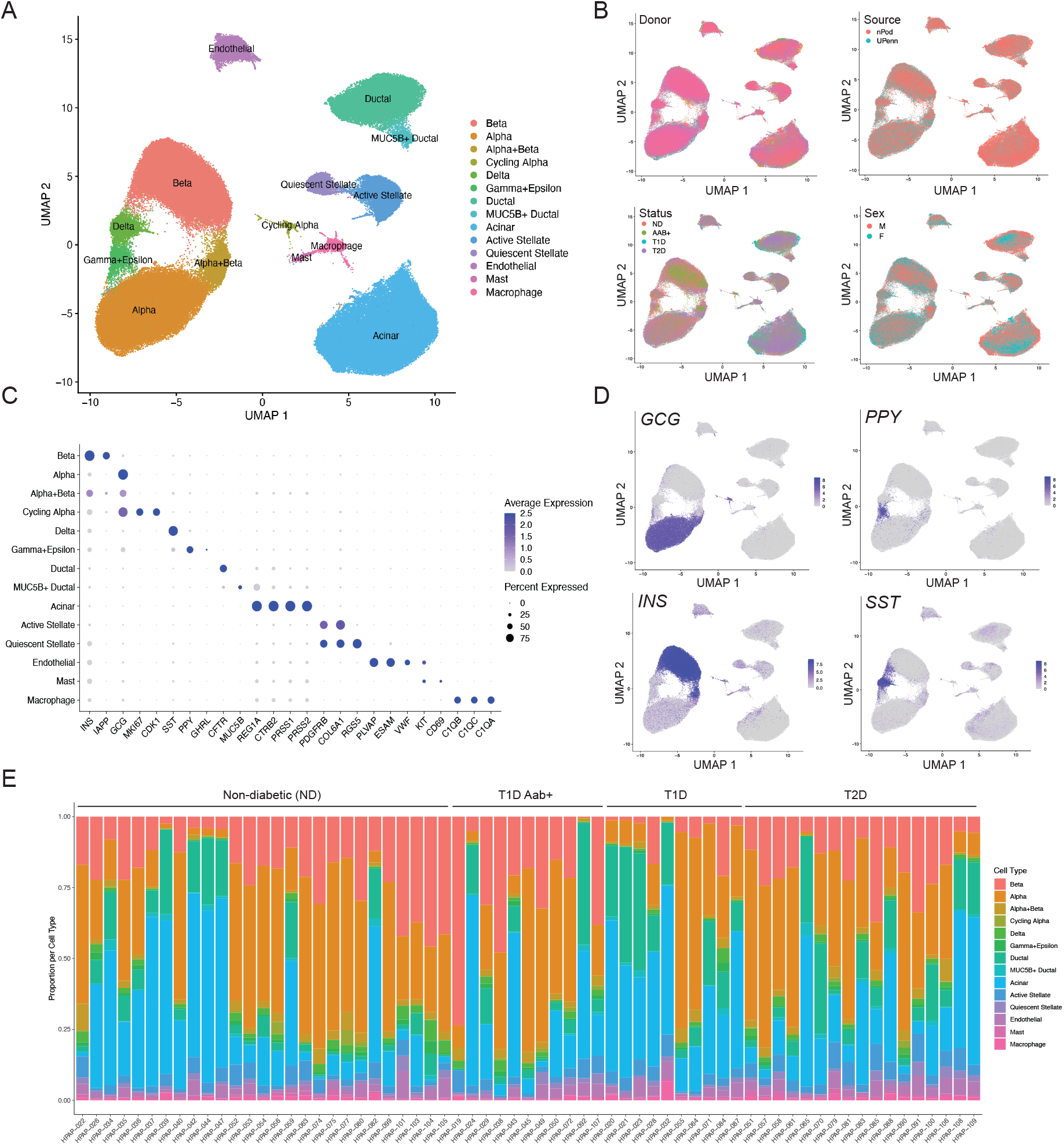
Map of gene expression in pancreatic islet cell types. (A) UMAP plot showing clustering of single cell RNA-seq profiles from 192,203 cells from HPAP donors in the PANC-DB website. Clusters are labeled based on cell type and sub-type identity using known marker genes. (B) Cells labeled based on variables such as donor, sex, disease status, and tissue source. (C) Dot plot showing normalized expression level and percent expressing cells for selected marker genes in each cluster. (D) Cells labeled with expression level of islet cell type hormones insulin (*INS*), glucagon (*GCG*), somatostatin (*SST*), and pancreatic polypeptide (*PPY*). (E) Proportion of cells generated from each donor from each cell type, grouped by disease state.

We next annotated the identity of clusters using a curated set of well-established cell type and marker genes (**Supplementary Table 2**). This revealed 10 total cell types including endocrine alpha (*GCG*), beta (*INS*), delta (*SST*), and gamma (*PPY*) cells as well as non-endocrine acinar (*REG1A*), ductal (*CFTR*), endothelial (*PLVAP*), stellate (*PDGFRA*), macrophage (*C1QA/B/C*), and mast cells (*KIT, CD69*) (**Figure 1C,D**). We also observed evidence for epsilon cells (*GHRL*) within the gamma cell cluster (**Figure 1C**), so we labeled this cluster as gamma+epsilon cells. Although islet cell type identity can be annotated using a small number of marker genes, knowledge of a larger set of genes specifically expressed in each cell type can provide potential additional insight into what drives cell identity. We therefore identified genes in each cell type with highly specific expression relative to other cell types in the study (**see Methods**). There were 542 genes with highly selective expression in one or more cell types (**Supplementary Table 3**), including those with no known function in the cell type. For example, in addition to canonical markers *INS* and *IAPP*, we identified beta cell-specific genes with known function in beta cells such as *RBP4* (18), *NPY* (19), *HADH* (20,21), and *C1QL1* (22) as well as those with no known beta cell function to our knowledge such as *SHISAL2B*, which could be targeted in future studies.

We also identified several cell types with multiple distinct clusters (**Figure 1A**). In most cases, these clusters represent previously described cell sub-types or states; for example, we identified quiescent and activated states of stellate cells (23), a MUC5B+ sub-population of ductal cells (23), and a sub-population of ‘cycling’ alpha cells (24) (**Figure 1A,C**). We identified 141 genes highly specific to a cell sub-type when compared to other cells of that same cell type (**Supplementary Table 4, see Methods**). We also identified a cluster comprised of both alpha and beta cells, which did not appear to be doublets, as the cells expressing insulin and glucagon were largely distinct (**Supplementary Figure 2**). These cells may represent cellular states of alpha and beta cells, as has been observed in other studies (15,24,25); however, due to the ambiguity over what cell populations these cells exactly represent, we excluded the cluster from downstream analyses.

We next compared the proportions of each islet cell type across samples (**Figure 1E**). As the purity of the islet preparations in PANC-DB varies dramatically, we assessed the proportion of different endocrine cell types as a function of the total number of endocrine cells per sample. Among non-diabetic samples, there was substantial variability in the proportion of different islet cell types; for example, the proportion of beta cells in islets ranged from .19 to .71. When considering disease states, we observed decreased proportion of beta cells in islets in T1D compared to ND control as expected (avg. ND=.40, avg. T1D=.18; Wilcox P=4.1×10^−4^). For beta cells, we observed a slight decrease in T2D (avg. ND=.40, avg. T2D=.365; Wilcox P=.36) and increase in T1D Aab+ (avg. ND=.40, avg. T2D=.49; Wilcox P=.24), although these estimates were not significant. While we also observed increased alpha cell proportion in T1D and T2D, this is likely explained by the relative decrease in beta cells.

We finally annotated the repertoire of genes expressed in each islet cell type in the non-diabetic state. For each cell type, we aggregated reads from all cells in non-diabetic samples, calculated normalized transcripts-per-million (TPM) expression levels from the ‘pseudo’-bulk counts for each gene, and defined genes expressed in a cell type (average TPM>1). There were between 7.6k-11.7k genes expressed per cell type in non-diabetic samples, with acinar cells (7.6k) and beta cells (8.2k) and having the smallest number of expressed genes. We next calculated gene expression levels for samples separately for each disease state. There were between 8.9k-11.9k genes expressed per cell type in T1D, 7.6k-11.7k genes expressed per cell type in T2D, and 7.4k-11.5k per cell type in T1D Aab+. Interestingly, for almost every cell type, we observed a larger number of expressed genes in both T1D and T2D compared to non-diabetic samples.

### Changes in islet cell type-specific gene expression in T1D and T2D

Identifying genes with changes in cell type activity in disease and pre-disease states can provide insight into disease pathogenesis. We therefore next determined changes in cell type gene expression in T2D (n=17), T1D (n=10), and T1D autoantibody positive (Aab+) non-diabetic (n=11) compared to non-diabetic control (n=27). We tested for differential expression of genes in a cell type from ‘pseudo’-bulk profiles across samples accounting for donor and technical characteristics as well as ambient background RNA signal (**see Methods**). In total, across all conditions, we identified 1,701 genes with significant changes in expression (FDR<.10) in at least one cell type (**Figure 2A**).

**Figure 2.**
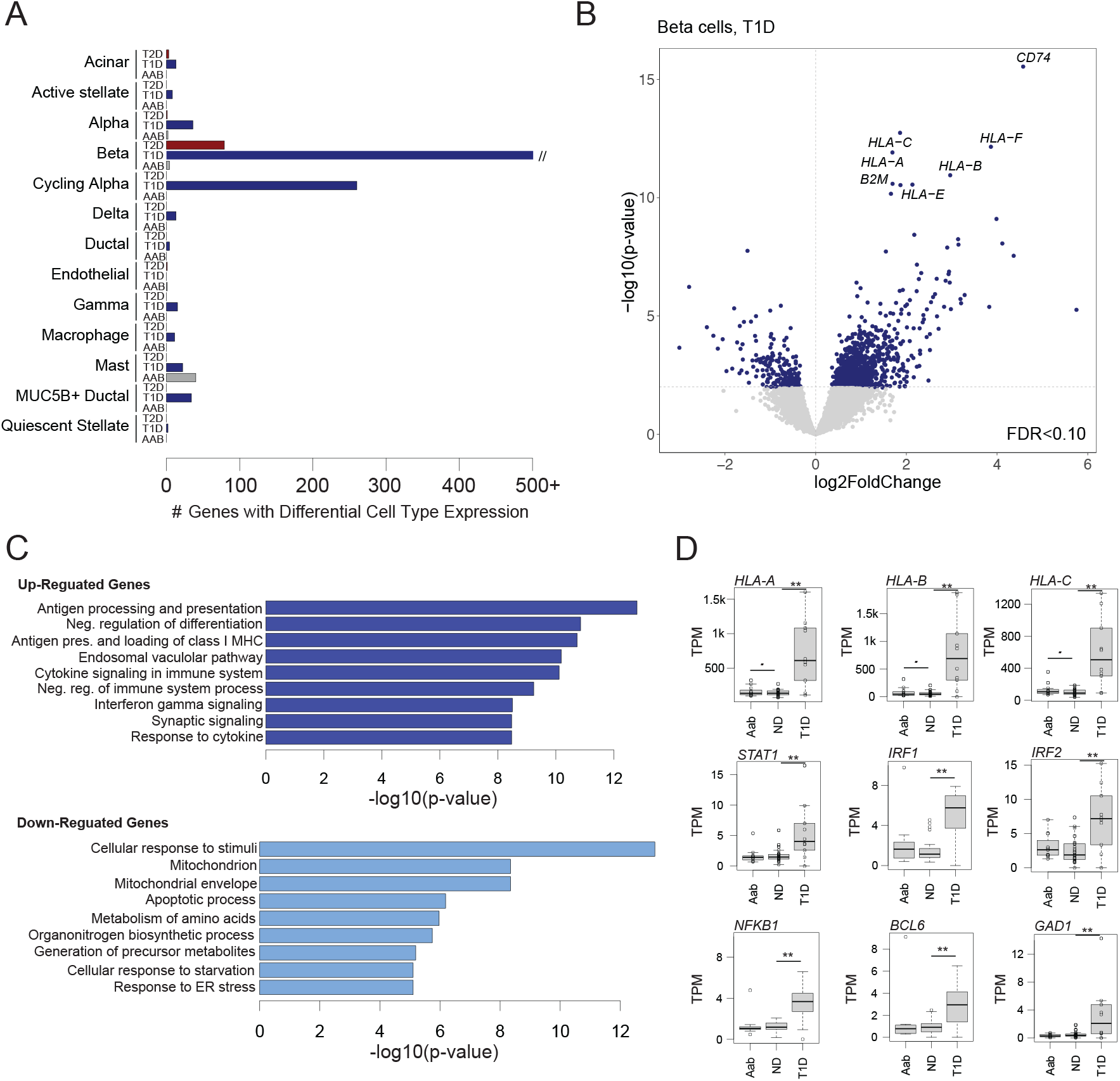
Changes in pancreatic islet cell type expression in disease states. (A) Number of genes in each cell type with significant changes (FDR<.10) in expression in T1D, T2D, and T1D autoantibody positive (AAB) compared to non-diabetic (ND) donors using DESeq2. The x-axis is capped at 500 genes for legibility, although the number of genes significant in beta cells in T1D is much higher. (B) Differential gene expression in beta cells in T1D. Genes with up-regulated expression are on the right side of the dashed line, and genes with down-regulated expression are on the left side. The hard line represents the p-value threshold corresponding to FDR<.10. Genes with the most significant increase in expression are labelled. (C) Gene sets enriched among genes with up-regulated and down-regulated beta cell expression in T1D. (D) Normalized beta cell expression level of HLA class I and selected stress response genes across ND, AAB and T1D donors. **FDR<.10, *uncorrected p<.10

For T2D, there were 84 genes with significant changes in any cell type compared to non-diabetic (ND) (**Supplementary Table 5**). Most of these genes were significant in beta cells (n=79), although we also observed a few differential genes in acinar, alpha, and endothelial cells. In beta cells, the genes with the largest increases in expression in T2D included *TSHR* which is the receptor for thyroid stimulating hormone, *SLC4A4* which is a bicarbonate co-transporter, and *TNFRSF11B* which is a cytokine receptor for tumor necrosis factor (TNF) family proteins. Genes with up-regulated expression in beta cells were enriched (FDR<.20) for biological processes related to protein modification, kinase activity, and ion binding, and by comparison down-regulated genes were enriched for processes related to protein transport and Golgi apparatus.

For T1D, we identified 1,808 genes with significant changes in expression compared to control (**Supplementary Table 6**). We observed the largest number of gene expression changes in beta cells (n=1,305), although there were also significant changes in cycling alpha cells (n=260), alpha cells (n=35), acinar cells (n=14), mast cells (n=22) and multiple other cell types. In beta cells, the genes with the largest increases in expression in T1D included multiple MHC class I genes (e.g. *HLA-A, HLA-B*) as well as genes involved in MHC processing and presentation such as *B2M* and *CD74* (**Figure 2B**). Genes up-regulated in beta cells were broadly enriched (FDR<.20) for biological processes related to MHC antigen processing and presentation (P=2.21×10^−17^), cytokine signaling (P=5.77×10^−14^) and differentiation (P=7.30×10^−15^), whereas down-regulated genes were enriched for processes related to stimulus response (P=5.19×10^−18^), response to ER stress (P=5.89×10^−9^), mitochondrial organization (P=1.02×10^−8^), response to metal ions (P=2.67×10^−8^), and apoptotic signaling (P=4.15×10^−8^) (**Figure 2C**). In addition to MHC class I genes, we observed up-regulation of genes mediating cytokine signaling responses such as *IRF1/2, STAT1/4*, and *NFKB1*, genes involved in cell survival such as *BCL6*, and the beta cell autoantigen *GAD1* (**Figure 2D**).

Finally, among individuals positive for T1D autoantibodies (Aab+) we identified very few genes with significant changes in expression compared to ND donors with no autoantibodies in any cell type (**Figure 2A**). While there was some evidence for up-regulation of MHC class I genes, we observed no clear change in expression of cytokine responsive factors such as IRFs, STATs and *NFKB1* (**Figure 2D**). By comparison, a recent study using largely the same set of samples from HPAP reported substantial changes in expression in beta cells in T1D Aab+ (28). Given the dramatic differences in the changes in expression between studies, more careful consideration of analytical approaches as well as larger sample sizes will be needed to reconcile these differences.

### Reference map and resource availability

The integrated map of gene expression in islet cell types generated by this study can be used to understand gene activity in physiological and disease states. In addition, this map can be used as part of bioinformatics pipelines, for example to perform reference mapping of new single cell RNA-seq datasets. We therefore provide several resources and interactive applications to facilitate the wide use of this integrated map in a variety of downstream analyses. These resources are all available at: http://www.isletgenomics.org.

First, we provide the islet cell type expression map in two interactive single cell browsers, CELLxGENE (29) and ShinyCell (30), which enable visualizing patterns across individual cells for example cell type identity, variables such as donor or library, technical factors such as number of features or percent mitochondrial reads, or the expression level of selected genes (**Supplementary Figure 3**). In addition, we provide the expression map within Azimuth (31), which can be used for rapid on-the-fly reference mapping of new datasets (**Supplementary Figure 4**).

Second, we provide annotations of activity in each islet cell type including marker genes, normalized gene expression levels, and changes in gene expression in T1D, T1D Aab+ and T2D. We developed several interactive applications that enable users to select specific genes to view expression levels in each cell type, as well as to visualize changes in cell type expression in different disease states (**Supplementary Figure 5**).

Finally, the analytical pipelines used for data processing and clustering, defining expressed genes in each cell type, and defining differentially expressed genes are provided open access.

## Discussion

Maps of gene expression levels in individual cell types within a heterogeneous tissue are valuable tools for hypothesis generation to understand cell type function and identity, gene activity, and changes in disease. In addition, these maps can be used for reference mapping of single cell RNA-seq datasets to facilitate annotation of cell identity and perform integrated analyses (31–33). While repositories such as PANC-DB provide access to a rich resource of raw sequence data and phenotypic information on human islet donors generated by HPAP (16), drawing insight from these data is a major challenge to researchers without single cell data analysis expertise. Our study provides an integrated map of gene expression profiles in islet cell types and changes in disease derived from the single cell RNA-seq experiments in HPAP, which will help enable downstream analyses and hypothesis generation for many non-single cell expert investigators.

There are several areas where the map can be further improved in future iterations. First, we were unable to separate a population of epsilon cells, likely due both to the rarity of epsilon cells and the sparsity of single cell RNA-seq profiles. By comparison, several studies profiling islets using different single cell technology resolved small epsilon cell populations (9,34). We also did not identify other rare cell types in the pancreas such as Schwann cells or lymphoid cell types. The samples profiled by HPAP are purified islets where other cell types outside of the islet microenvironment are preferentially removed, and therefore the profiles of non-islet cell types are under-represented. Furthermore, the repertoire of discrete states that exist within each cell type, as well as any sub-types for example with distinct spatial localizations, remains to be resolved. Continued profiling of donors and cells from both purified islets and whole pancreas will help to define profiles for all pancreatic cell type and sub-types. Finally, even after accounting for ambient background RNA there is still residual expression of genes in off-target cell types, particularly for highly expressed genes in common cell types, where improvements in background correction methods for single cell RNA-seq are needed to estimate islet cell type-specific expression more accurately.

Genes with significant changes in cell type-specific expression in T1D and T2D provide insight into disease pathogenesis, for example up-regulation of MHC class I genes and cytokine signaling processes in T1D and differentiation related processes in T2D. There were few changes in gene expression in T1D Aab+ where we identified almost no significant genes. By comparison, a recent study using largely the same set of HPAP samples found marked changes in beta cell expression profiles in T1D Aab+ (28). Given substantial heterogeneity between donors and in disease processes, much larger sample sizes will be needed to determine the true extent to which gene profiles change in different disease and physiological states. Larger sample sizes will also enable finer grained partitioning of samples to understand gene regulatory differences between phenotypic sub-groupings. For example, many Aab+ samples in HPAP are single GAD+ (28), yet there is great diversity in the number and type of autoantibodies that individuals can have with different rates of progression to T1D (35,36). Similarly, profiling islets from impaired glucose tolerant donors will help understand changes that occur during progression to T2D (37).

In summary, our map of islet cell type-specific expression and associated resources of cell typespecific gene activity in physiological and disease states provided by this study will be a valuable reference to the islet and diabetes research community.

## Methods

### Data Availability

Organ procurement and processing was performed by the Human Pancreas Analysis Program (HPAP) as previously described (16). Single cell RNA-sequencing (scRNA-seq) data from isolated and dissociated pancreatic islets were made publicly available by HPAP and raw fastq files for experiments from 67 donors (10 T1D, 17 T2D, 29 ND, 11 ND but AAB+) were downloaded from the PANC-DB data portal. Cell Ranger 6.0.1 (10x Genomics) software was used to perform alignment to the GRCh38 human reference genome and generate count matrices.

### Preliminary filtering

Barcodes were filtered for a minimum of 500 expressed genes per cell and less than 15% mitochondrial reads. Two samples (HPAP-027 and HPAP-093) were removed since the mean number of expressed genes per cell after this filtering step was markedly lower than for other samples (<1000).

### Ambient RNA correction

Ambient RNA removal was performed to account for extracellular RNA contamination that may get trapped in a droplet during library generation. SoupX 1.6.1 (38) was used on raw feature barcode matrices for ambient RNA removal on the remaining 65 samples using the automated contamination fraction estimation method. Raw count values for each sample were corrected using the SoupX contamination estimates and the round to integer feature, ensuring resulting counts remain integers for use in downstream analyses (39).

### Data processing and clustering

The SoupX-corrected count matrices were merged and log normalized with a scale factor of 1000. The variance stabilizing transformation (vst) method was used to find the 2000 most variable features. Data was scaled and principal component analysis was performed with 20 principal components using Seurat 4.2.0 (31). Harmony 0.1.1 (40) was used for batch correction using donor, 10x Genomics assay chemistry (10x 3’ v2 or 10 3’ v3), and tissue source (nPOD or UPenn) as covariates. Uniform manifold approximation and projection (UMAP) and neighbors were calculated using the reduction from Harmony. Clustering was performed in Seurat 4.2.0 using the Leiden algorithm at a resolution of 0.5.

### Post-clustering doublet removal

Scrublet 0.2.3 (41) was used to identify doublets with the default parameters (expected doublet rate of 6%, minimum counts of 2, minimum cells of 3, minimum gene variability percentile of 85%, and 30 principal components). For each sample, RNA count matrices were extracted, saved in MatrixMarket format, and input into Scrublet with default parameters. There were 4,382 barcodes flagged as doublets, and we removed these barcodes from the merged Seurat object and re-performed Harmony integration and clustering (resolution 0.3) with the remaining barcodes as described above. We further curated a set of cell type-specific marker genes, and clusters that contained marker genes for two or more different cell types were further sub-clustered using the Leiden algorithm at resolutions of 0.15-0.25. Any sub-clusters expressing multiple cell type marker genes were presumed to be residual doublets, and we manually removed these sub-clusters representing a total of 13,036 barcodes. We then re-performed Harmony integration and clustering using the final set of barcodes as described above.

### Cell type specific marker genes

While there are well established marker genes for many pancreatic islet cell types, we performed an unbiased analysis of cluster-specific marker genes using the FindAllMarkers function in Seurat 4.2.0 (31). For each cluster, a Wilcoxon rank sum test was run on log-normalized counts in comparison to the remaining clusters. Genes were considered markers if they were expressed in more than 25% of cells in the cell type and less than 25% of other cells, had log2 fold change threshold greater than 1, and had adjusted P-value less than .05. For cell types with multiple clusters, we used the FindMarkers function to identify genes with cluster-specific activity relative to other cells in the same cell type.

### Cell type gene expression profiles

We aggregated reads from cells in each cell type and created ‘pseudo’-bulk counts from contamination-corrected RNA counts. We calculated transcripts per million (TPMs) for each donor in each cell type using GENCODE v38 GRCh38.p13 (42) gene size annotations. Differential gene expression analyses were performed using DESeq 1.34.0 (43), comparing cell type profiles between ND controls and T1D Aab+, T1D, and T2D. Sex, scaled age, scaled BMI, 10x kit chemistry, and tissue procurement source were included as covariates. Genes were only tested for a cell type if at least half of the samples per tested condition had at least 5 counts. Multiple test correction was performed using the Benjamini-Hochberg false detection rate correction at an alpha of 10%. We performed gene set enrichment analyses (GSEA) (44) for up- and down-regulated genes in beta cells using Gene Ontology, KEGG pathway, and REACTOME terms. For conditions with less than 100 genes differentially expressed at FDR<.10, we used an uncorrected p-value threshold of .001 as input to GSEA. From the results of GSEA we considered terms significant at FDR<.20 and which also contained less than 1000 genes.

## Data availability

The raw sequence data is available on the PANC-DB website. Processed files and derived annotations generated by this study are available at isletgenomics.org. Custom code is available at https://github.com/Gaulton-Lab/HPAP-scRNA-seq.

## Author contributions

K.J.G. conceived of the study and obtained funding. K.J.G. and R.M.E, wrote the manuscript. R.M.E. performed all single cell data analyses. R.M.E., R.L.M., H.M.M., P.B., and ML.O. developed analytical pipelines for single cell data processing and analysis. P.K. developed and implemented tools for data visualization and interaction.

## Acknowledgements

This work was supported by the National Institutes of Health grant numbers DK-105554, DK-114650, and DK-120429 to KJG. This manuscript used data acquired from the Human Pancreas Analysis Program (HPAP-RRID:SCR_016202) Database, a Human Islet Research Network (RRID:SCR_014393) consortium (UC4-DK-112217, U01-DK-123594, UC4-DK-112232, and U01-DK-123716).

---

**Supplementary Figure 1.**
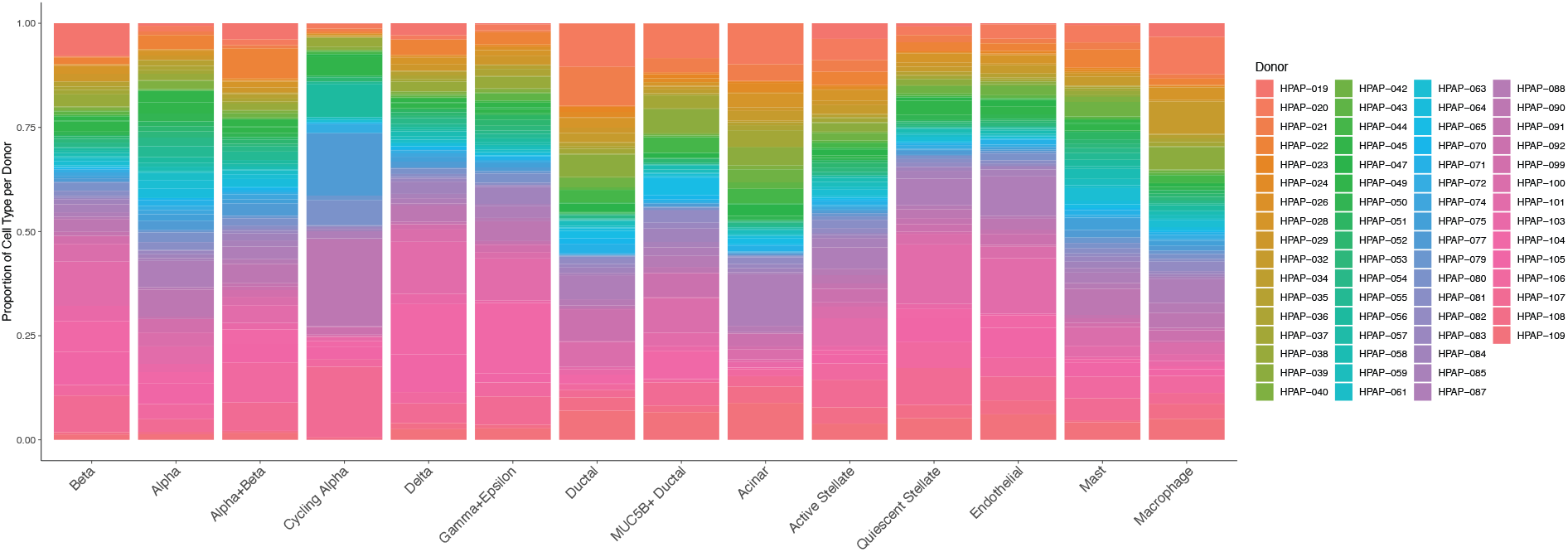
Sample representation per cluster. Proportion of cells in each cluster from all donor samples in HPAP. No clusters have representation from only a small number of donors.

**Supplementary Figure 2.**
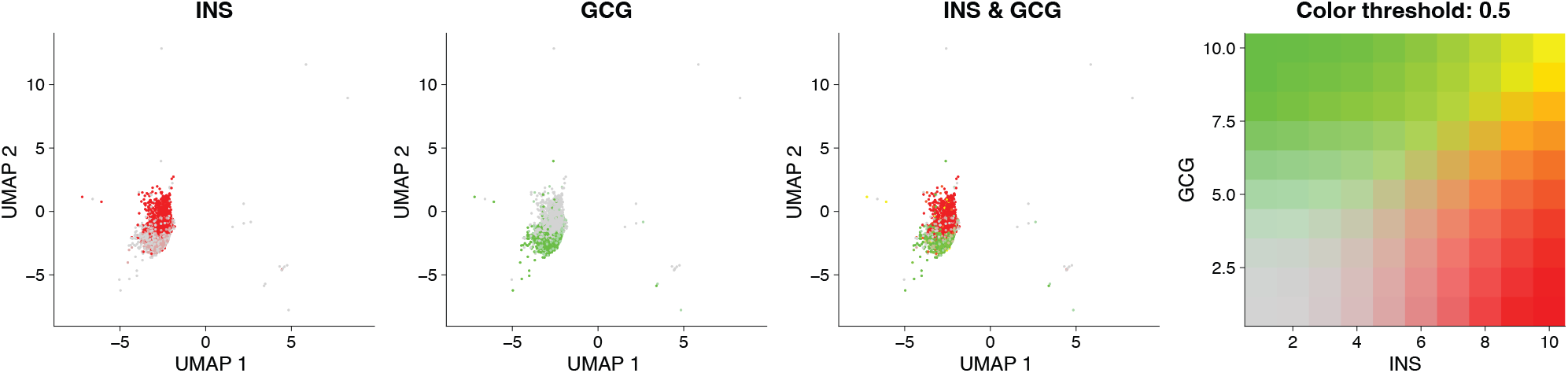
Marker gene expression in alpha+beta cluster. Expression level of insulin and glucagon separately, as well as overlaid together, in the alpha+beta cluster. The insulin and glucagon expressing cells within this cluster appear largely distinct.

**Supplementary Figure 3.**
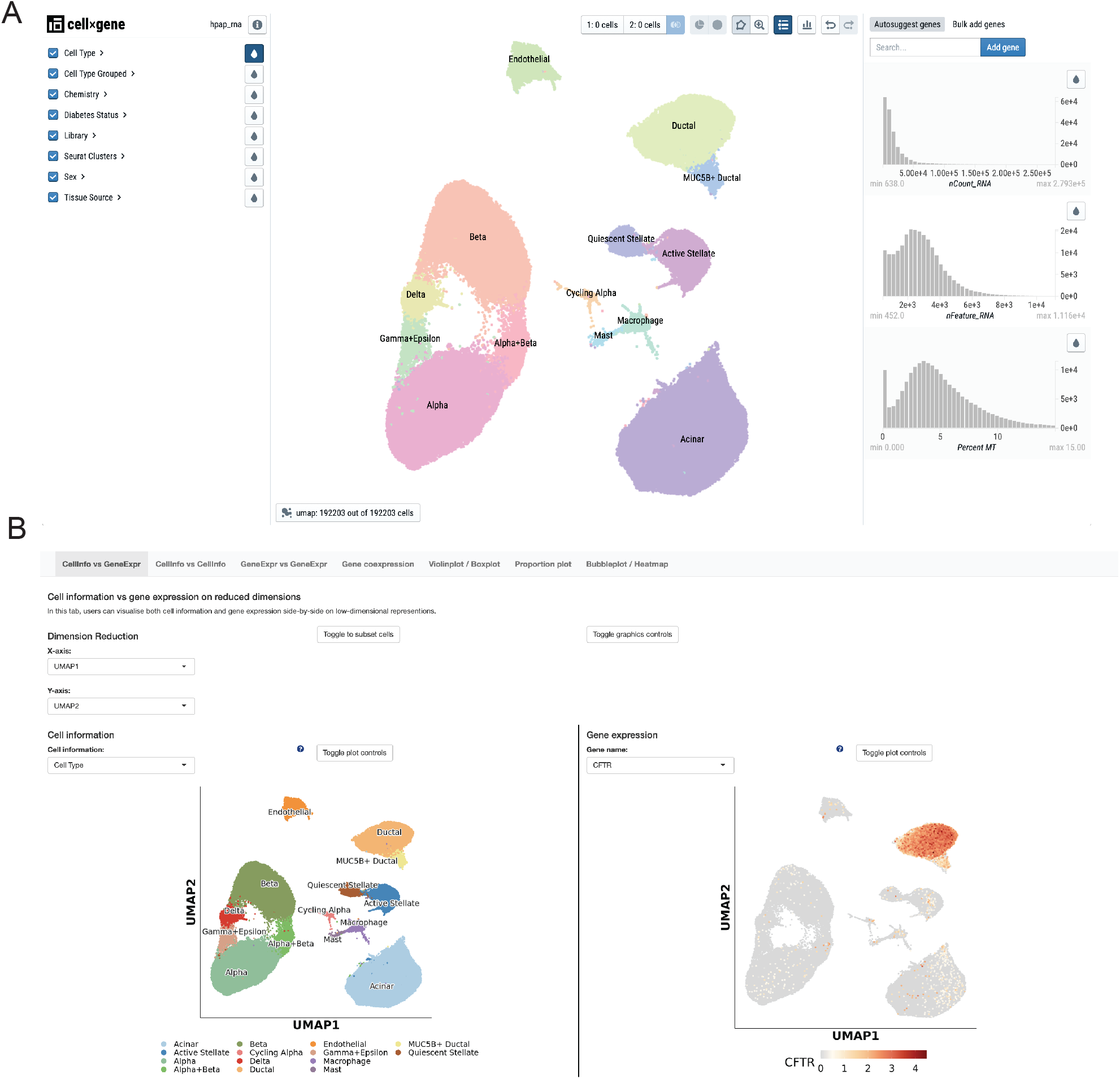
Visualizing single cell profiles in islet cell types. (A) Islet single cell object in Cellxgene browser including cell type labels and technical features. (B) Islet single cell object in ShinyCell including cell type labels and expression of selected gene.

**Supplementary Figure 4.**
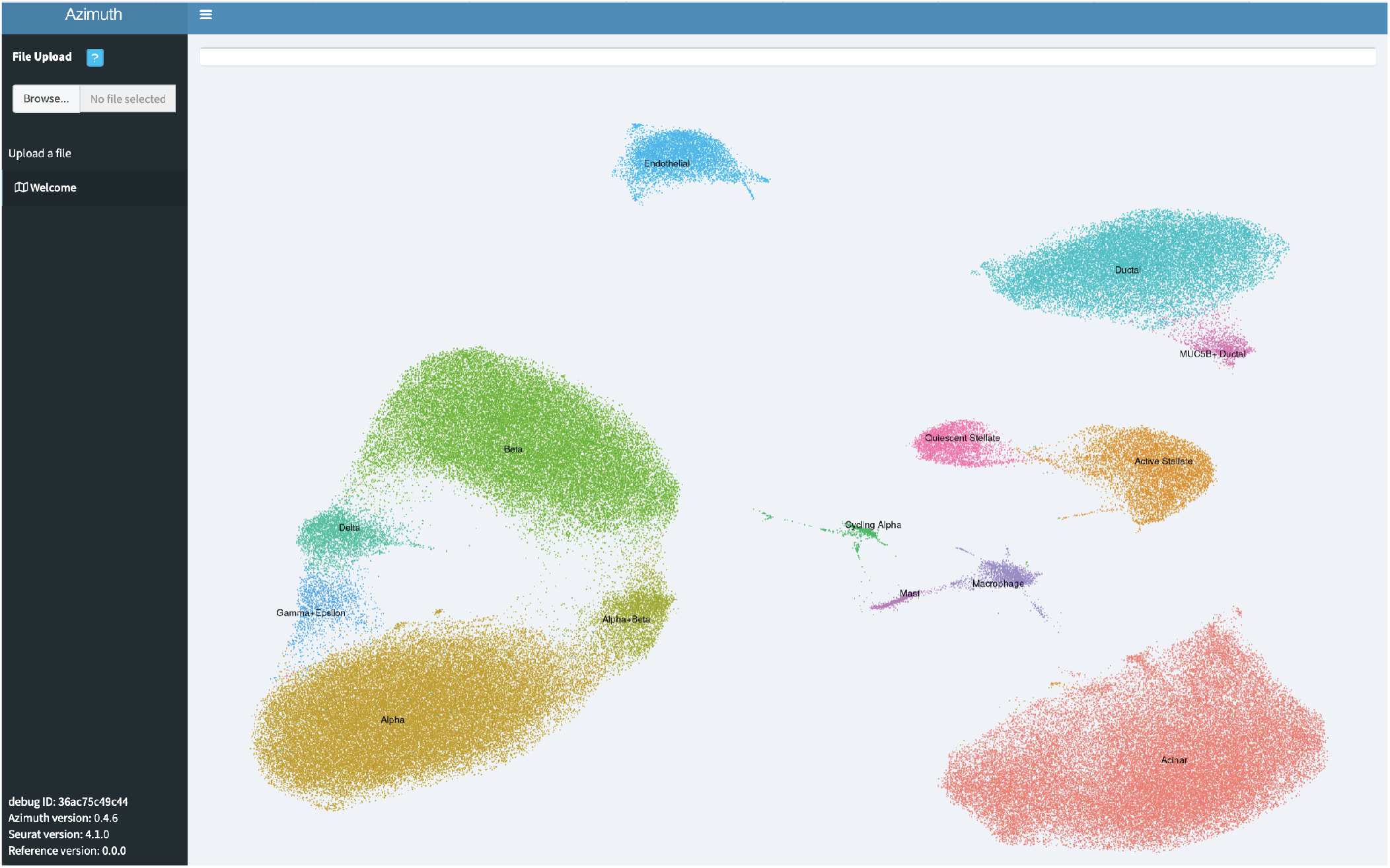
Reference mapping of islet single cell gene expression. Screenshot of islet cell type expression in the Azimuth application which enables users to upload their own data and perform annotation of cell type identity.

**Supplementary Figure 5.**
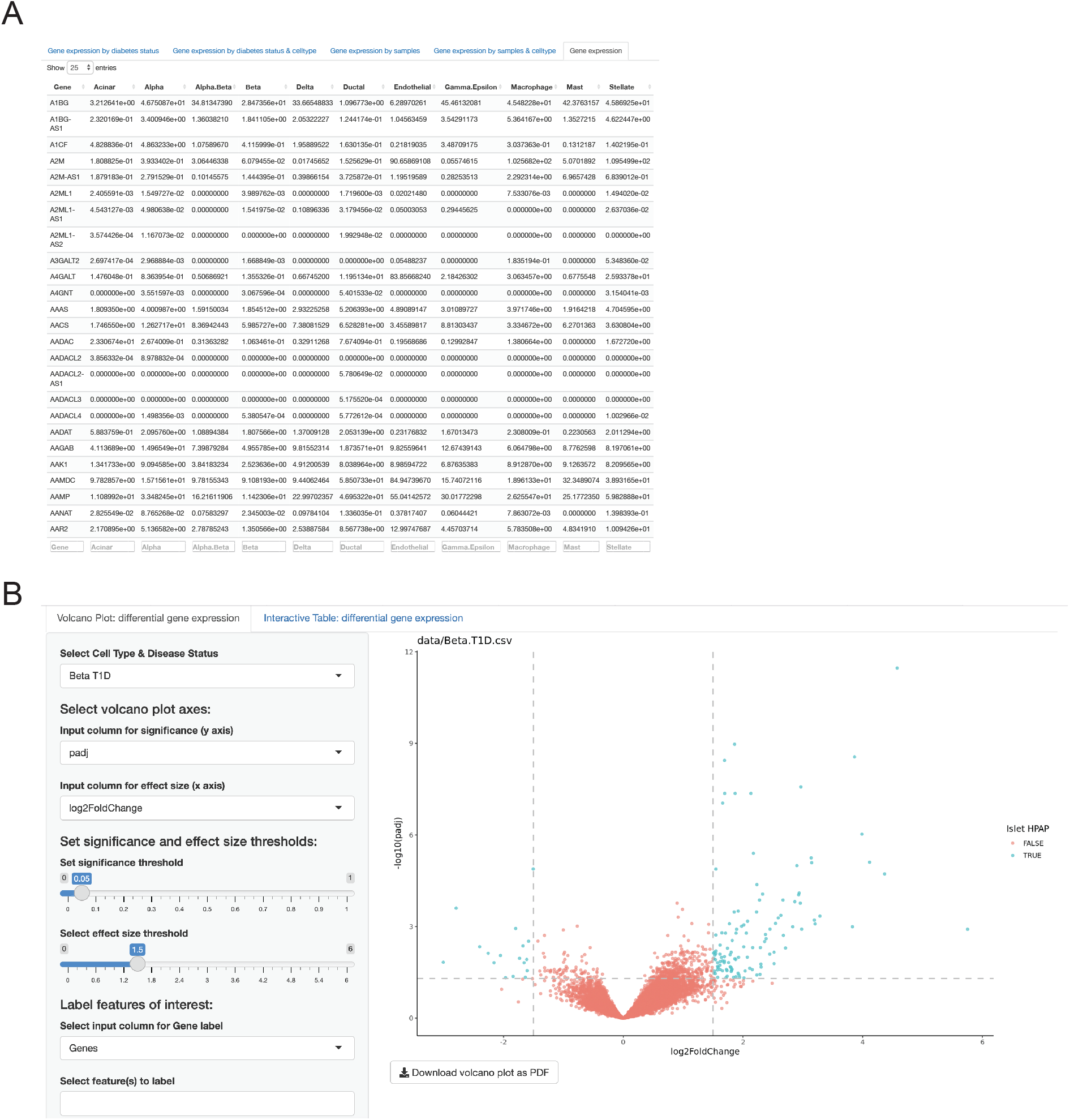
Querying and visualizing islet cell type-specific expression. (A) Interactive table showing normalized expression levels of genes in each identified cell type. (B) Interactive visulization of differential cell type expression in disease states.

